# Reversible Pulmonary Trunk Banding. XI: Myocardial Vascular Endothelial Growth Factor Expression in Young Goats Submitted to Ventricular Retraining

**DOI:** 10.1101/644278

**Authors:** Renato S. Assad, Eduardo A. V. Rocha, Vera D. Aiello, Tiago A. Meniconi, Maria C. D. Abduch, Petronio G. Thomaz, Marcelo B. Jatene, Luiz F. P. Moreira

**Author notes:** All the authors contributed equally to this work. **Corresponding Author:** Renato S. Assad, MD, PhD, Heart Institute University of São Paulo Medical School, Division of Surgical Research, Ave. Dr. Eneas C. Aguiar, 44, São Paulo, SP – Brazil 05403-000, /.

## Abstract

**Background:** Ventricle retraining has been extensively studied by our laboratory. Previous studies have demonstrated that intermittent overload causes a more efficient ventricular hypertrophy. The adaptive mechanisms involved in the ventricle retraining are not completely established. This study assessed vascular endothelial growth factor (VEGF) expression in the ventricles of goats submitted to systolic overload.

**Methods:** Twenty-one young goats were divided into 3 groups (7 animals each): control, 96-hour continuous systolic overload, and intermittent systolic overload (four 12-hour periods of systolic overload paired with 12-hour resting period). During the 96-hour protocol, systolic overload was adjusted to achieve a right ventricular (RV) / aortic pressure ratio of 0.7. Hemodynamic evaluations were performed daily before and after systolic overload. Echocardiograms were obtained preoperatively and at protocol end to measure cardiac masses thickness. At study end, the animals were killed for morphologic evaluation and immunohistochemical assessment of VEGF expression.

**Results:** RV-trained groups developed hypertrophy of RV and septal masses, confirmed by increased weight and thickness, as expected. In the study groups, there was a small but significantly increased water content of the RV and septum compared with those in the control group (p<0.002). VEGF expression in the RV myocardium was greater in the intermittent group (2.89% ± 0.41%) than in the continuous (1.80% ± 0.19%) and control (1.43% ± 0.18%) groups (p<0.023).

**Conclusions:** Intermittent systolic overload promotes greater upregulation of VEGF expression in the subpulmonary ventricle, an adaptation that provides a mechanism for increased myocardial perfusion during the rapid myocardial hypertrophy of young goats.

## 1 INTRODUCTION

The ideal surgical treatment for transposition of the great arteries (TGA) is the Jatene procedure performed during the neonatal period. Regarding late referrals to tertiary centers in developing countries, rapid ventricular retraining for completion of anatomic correction in 2 stages still remains as an option. [1] Nevertheless, the performance of the trained ventricle may not be ideal. [2] In the long run, some degree of ventricular dysfunction may occur, with a consequent increase in late mortality. [3] Lim et al. have demonstrated that ventricular retraining is a risk factor after anatomic correction of TGA. [4] In the Boston series, late ventricular dysfunction occurred in almost 25% of the patients submitted to the 2-stage arterial switch operation. [5] Previous experimental studies about subpulmonary ventricular retraining have demonstrated increased areas of necrosis and/or fibrosis in hearts submitted to traditional pulmonary artery banding (PAB). [6] These morphologic observations might explain why patients who undergo the primary arterial switch operation have better long-term outcomes than those submitted to the 2-stage operation for TGA. [7] Therefore, the ideal subpulmonary ventricular retraining protocol remains controversial. Studies from our laboratory have demonstrated that it may be possible to equalize the right and left ventricular masses of young animals with only 96 hours of systolic overload of the right ventricle. [8] Intermittent systolic overload has been demonstrated to be superior to traditional PAB, resulting in a more efficacious ventricular hypertrophy with less exposure to pressure overload. Moreover, myocardial performance under pharmacological stress was better in the animals submitted to intermittent systolic overload compared to traditional PAB. [9] There is a great interest in the adaptive mechanisms of the retrained ventricle. Abduch et al. have demonstrated that cellular proliferation was similar between ventricles traditionally and intermittently trained. [10] Nevertheless, the rate of the myocardial vascular endothelial cells’ proliferation during the process of ventricular retraining remains unclear. Ideally, there should be an increase in the number of capillaries to support the increase in myocardial mass and systemic vascular resistance in the retrained ventricle. In the present study, we analyzed the expression of vascular endothelial growth factor (VEGF), which, together with other growth factors, plays a physiological role in regulating vascular development. VEGF is a mitogen factor for endothelial cells, and it selectively permeabilizes the endothelium and plasma proteins without causing injury. [11,12] These characteristics are essential for angiogenesis. Experimentally, VEGF has been used via gene therapy to optimize the healing of myocardial infarction, increasing the capillary density and reducing the size of the infarcted area. [13] This study aims to experimentally examine the adaptation of the subpulmonary ventricle with regard to induction of myocardial angiogenesis signaling in response to pressure overload, using an adjustable PAB system.

## 2 MATERIALS AND METHODS

Twenty-one healthy young male goats, aged between 30 and 60 days and of comparable weights (p=0.08), were randomly divided into 3 groups: control (n = 7; weight, 12.0 ± 1.0 kg), continuous (n = 7; weight, 9.4 ± 0.6 kg), and intermittent (n = 7; weight, 9.8 ± 0.8 kg). All animals received humane care in compliance with the ARRIVE guidelines established by the Guide for the Care and Use of Laboratory Animals (The National Centre for the Replacement, Refinement and Reduction of Animals in Research, London, UK, 2010). All animals experiments were carried out in accordance with the U.K. Animals (Scientific Procedures) Act, 1986 and associated guidelines, EU Directive 2010/63/EU for animal experiments. The protocol (#093/14) was reviewed and approved by the Institutional Ethics Committee for Research Protocols.

### 2.1 ANESTHESIA

Anesthesia was conducted as previously reported. [9] Each animal was monitored with an electrocardiogram (ECG) and prepared for a sterile procedure. Antibiotics (cefazolin 500 mg intravenously and gentamicin 40 mg intramuscularly) were administered daily during the study. All of the goats were extubated right after the surgical procedure and remained ambulatory and breathing spontaneously throughout the protocol.

### 2.2 OPERATIVE TECHNIQUE

All operations were performed by a single surgeon through a left lateral thoracotomy. The lung was retracted laterally to allow exposure of the pericardial sac and descending aorta. A 17-gauge heparinized catheter was inserted in the right ventricular (RV) outflow tract, main pulmonary artery (PA), and descending aorta (Ao) for pressure measurements at specific time intervals throughout the protocol. Pressure measurements were taken with computer software (ACQknowledge, version 3.01; Biopac Systems, Inc, Goleta, Calif). After every pressure measurement, the catheter was filled with heparin. In addition, heparin (5,000 UI) was given subcutaneously daily throughout the study period to help maintain catheter patency. The pulmonary trunk was dissected free for adjustable banding device implantation (SILIMED – Silicone e Instrumental Médico-Cirúrgico e Hospitalar Ltda., Rio de Janeiro, RJ, Brazil), as previously reported. [14]

### 2.3 RV SYSTOLIC OVERLOAD PROTOCOL

Baseline hemodynamic data (RV, PA, and Ao pressures) were collected by a single researcher in a conscious animal with the adjustable banding system deflated. RV training was begun after a 72-hour convalescence period with percutaneous PAB insufflation with saline solution to achieve an RV/systemic pressure ratio of 0.7, limited by a 10% drop in systemic systolic blood pressure. Pressure measurements in the Ao, right ventricle and PA were taken daily in the 3 groups.

#### 2.3.1 Continuous Group

The PAB device was readjusted every morning throughout the protocol to keep the continuous RV/aortic pressure ratio at 0.7 during the study period of 96 hours. Hemodynamic data were collected before and after PAB readjustments.

#### 2.3.2 Intermittent Group

The goats underwent 4 daytime periods of 12-hour systolic overload, alternating with a 12-hour nighttime resting period, during the same period of 96 hours as the continuous group. Hemodynamic data were collected twice a day (every 12 hours) during PAB readjustments.

#### 2.3.3 Control Group

The PAB system was maintained deflated during the entire protocol. Hemodynamic data were collected once daily (mornings).

### 2.4 ECHOCARDIOGRAPHY

A single experienced observer conducted all examinations blinded to study group with the animals under light sedation (ketamine 15 mg intramuscularly), prior to the beginning and at the end of the protocol. Image acquisition was obtained through 7.5 MHz and 2.5 MHz multifrequency transducers (Acuson Cypress, Siemens, Erlangen, Germany). The end-diastolic thicknesses of the cardiac masses were measured by 2-dimensional echocardiography through the parasternal long-axis view, at the level of the mitral valve leaflet tips. [15]

### 2.5 MORPHOLOGY

The animals were killed after 96 hours of the study protocol. The heart was harvested and cardiac samples were drawn from the right ventricle, left ventricle, and ventricular septum to be subsequently analyzed for VEGF tissue expression.

#### 2.5.1 CARDIAC MASS WEIGHTS

The epicardial fat, both atria, and semilunar valves were carefully dissected from the heart; the right ventricle, left ventricle, and ventricular septum were separated and individually weighed (METTLER AE-200; Mettler-Toledo AG, Greifensee, Switzerland). Cardiac mass weights were indexed to body weight (g/kg).

#### 2.5.2 WATER CONTENTS OF THE MASSES

The water content was obtained individually in each cardiac chamber by subtracting the collected sample weight at autopsy from the weight of the dehydrated chamber (70 hours at 60° C). Values were obtained as a percentage of weight change, assuming that the water distribution was homogenous in the septum and in the ventricles.

#### 2.5.3 VEGF EXPRESSION

All of the hearts, including those from animals in the control group, were fixed in 10% buffered formalin for 24 hours. Transversal sections of both ventricles and the ventricular septum were obtained. After routine histologic processing, 5-μm sections underwent immunohistochemical staining for VEGF A-20 (primary antibody from Santa Cruz Biotechnology, Dallas, USA, dilution 1/1500), using the streptavidin-peroxidase method. [16] Sections were counterstained with Harris hematoxylin (Merck Millipore, Darmstadt, Germany). The slides were evaluated using an image analyzer (AxioVision, software V.4.7.1.0, Carl Zeiss AG). For quantification of the area fraction occupied by VEGF-labeled cells, 25 pictures of perivascular myocardial regions were obtained from each section of the heart (right ventricle, left ventricle, and ventricular septum) under objective magnification x40. The sum of the cytoplasmatic areas labeled by the antibody was quantified by color detection in each microscopic field and expressed as the area fraction (green color). Figure 1 shows a section of the myocardium from an animal of the Intermittent group. Notice the presence of green spots corresponding to the labeled cells after automatic color detection of VEGF labeling in cells of the vessel walls and perivascular region. For each section of the heart, the mean area occupied by the labeled cells was calculated.

**Figure 1.**
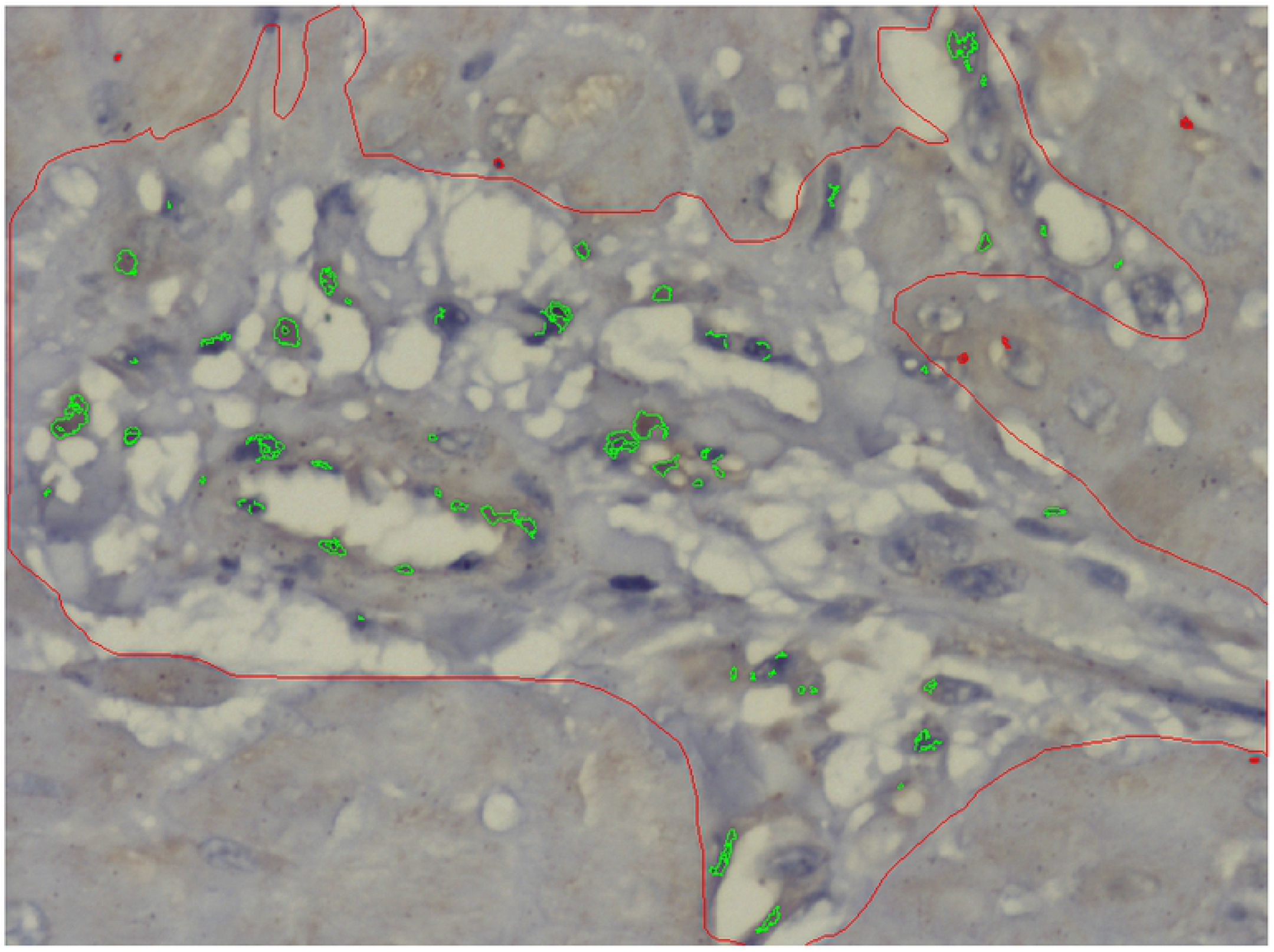
Photomicrograph of the myocardium from an animal of the Intermittent group, counterstained with Harris hematoxylin. The perivascular area is indicated manually (red line). Notice the presence of green spots corresponding to the labeled cells after automatic color detection of VEGF labeling in cells of the vessel walls and perivascular region. Objective magnification= 40X. Image Analyzer: AxioVision, software V.4.7.1.0, Carl Zeiss AG

### 2.6 STATISTICAL ANALYSIS

Data were analyzed using GraphPad Prism 5 software (GraphPad Inc, La Jolla, CA). Values are expressed as the mean ± SEM. Two-way analysis of variance (ANOVA) was used to compare groups and cardiac masses, followed by the Tukey post hoc test. For all tests, the significance level was 5%.

## 3 RESULTS

### 3.1 HEMODYNAMIC MEASUREMENTS

Both RV systolic overload protocols did not affect goat systemic hemodynamics. Figure 2 shows the RV-PA systolic pressure gradient throughout the protocol. It increased in both study groups (p<0.001) and in every time period (p<0.001). The PAB gradients in both study groups were significantly higher than their respective baseline values and the PAB gradient in the control group (P<0.001). At the end of the protocol, the banding gradient generated by the intermittent group was significantly higher than that generated by the continuous group (p<0.001).

**Figure 2.**
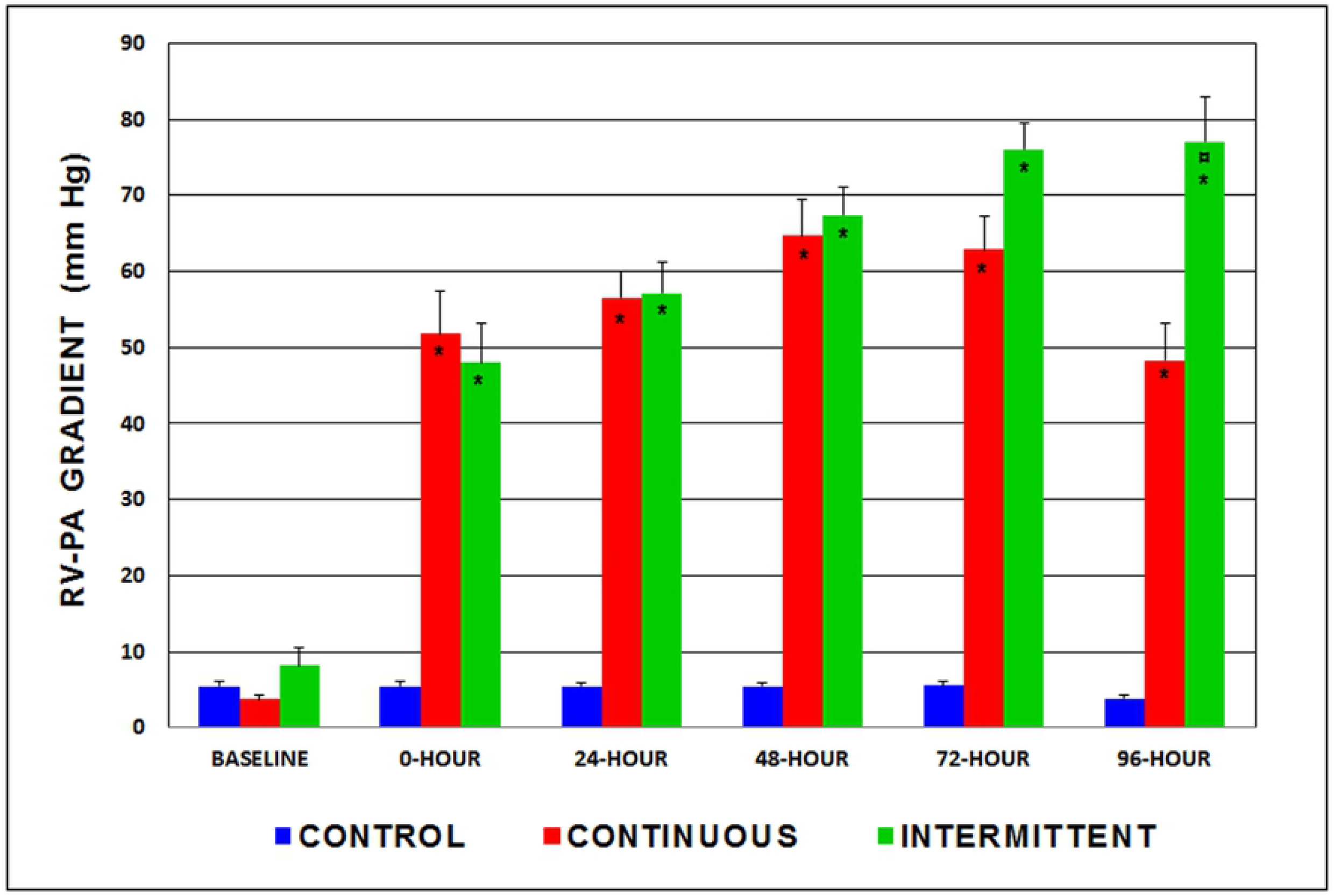
Right ventricular (RV) to pulmonary artery (PA) systolic pressure gradient (mm Hg) throughout the protocol. * p <0.001 when compared with respective baseline and control group values; **¤** p<0.001 when compared with 96-hour continuous group. n= 7 in each group.

### 3.2 ECHOCARDIOGRAPHIC FINDINGS

Table 1 shows the RV wall thicknesses of the 3 groups at the beginning and at the end of the systolic overload protocol. Compared with the control group and the baseline measurements, both study groups showed an increased thickness of the RV free wall at the end of the protocol (p<0.001), which was significantly greater in the intermittent group (129.2%) than the continuous group (58.2%; p <0.001). The thicknesses of the septal wall and left ventricle in the 2 study groups did not significantly differ from control values, thus, were not affected by the RV systolic overload protocol.

**Table 1.**
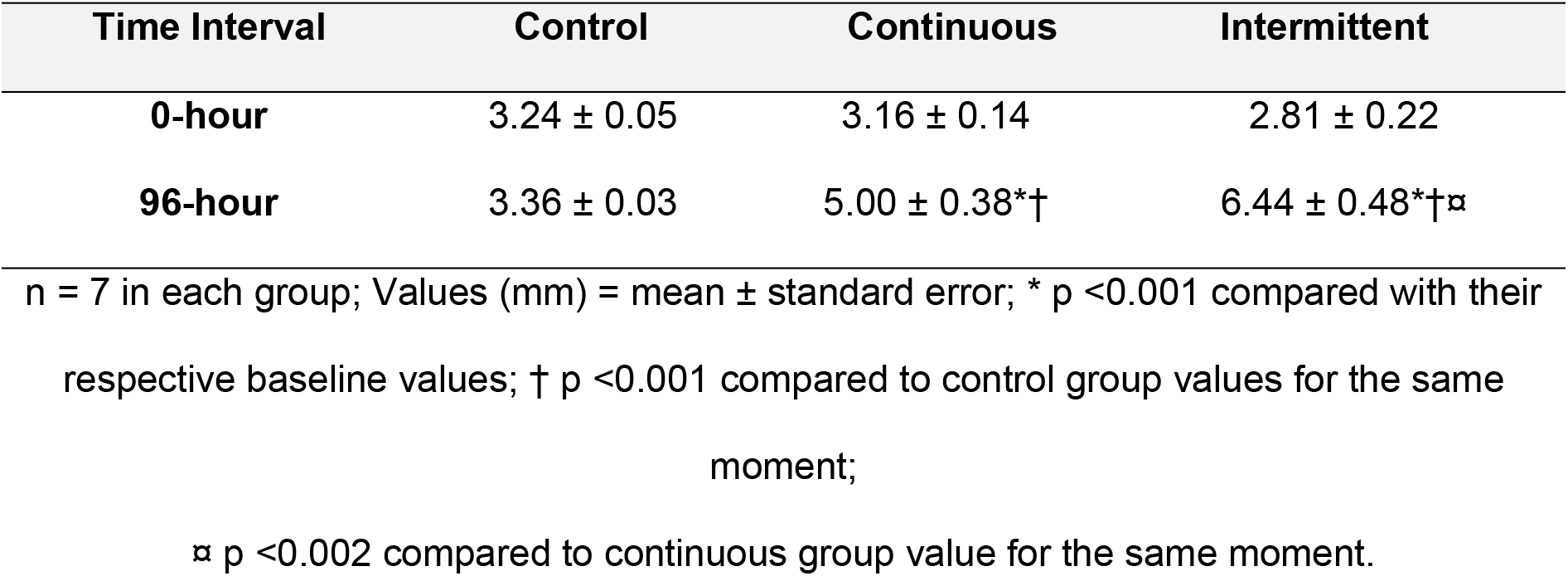
Right ventricular wall thicknesses in the control, continuous, and intermittent groups, measured by echocardiography.

| Time Interval | Control | Continuous | Intermittent |
| --- | --- | --- | --- |
| 0-hour | 3.24 ± 0.05 | 3.16 ± 0.14 | 2.81 ± 0.22 |
| 96-hour | 3.36 ± 0.03 | 5.00 ± 0.38*† | 6.44 ± 0.48*†¤ |

n = 7 in each group; Values (mm) = mean ± standard error; \* $p < 0.001$ compared with their respective baseline values; † $p < 0.001$ compared to control group values for the same moment;
¤ $p < 0.002$ compared to continuous group value for the same moment.

### 3.3 MORPHOLOGIC FINDINGS

#### 3.3.1 CARDIAC MASS WEIGHTS

Both study groups showed significant increases in RV mass (intermittent: 115.8%; continuous: 90.8%; p<0.001) and septal mass (intermittent: 55.8%; continuous: 45.4%; p<0.047) weights compared to those in the control group (Figure 3). LV mass was not altered by the RV training protocols.

**Figure 3.**
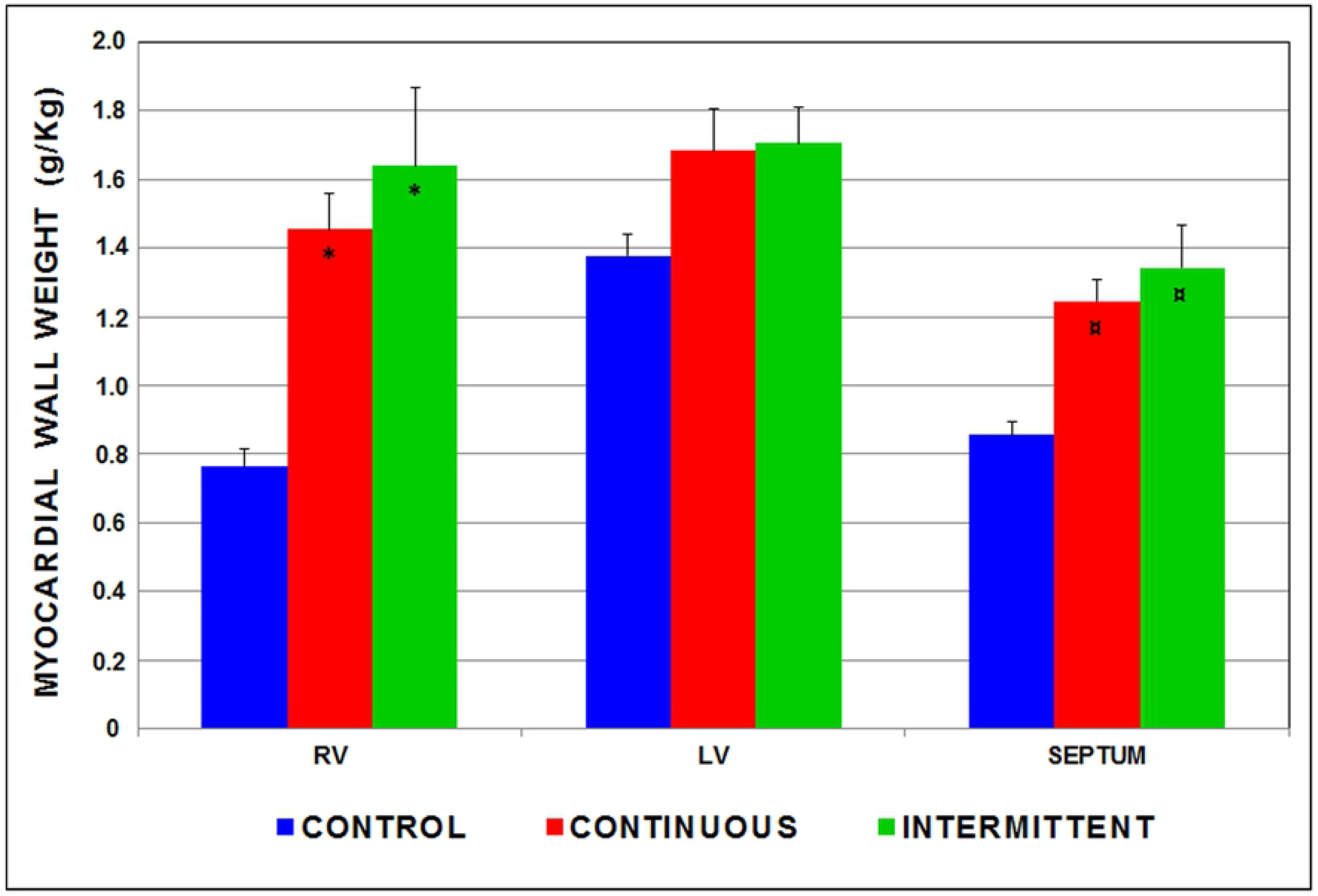
Weights for the right ventricle (RV), left ventricle (LV), and septum indexed to body weights of the 3 groups. * p<0.001 when compared to the control group; **¤** p<0.047 when compared to the control group. n= 7 in each group.

#### 3.3.2 WATER CONTENT

Table 2 shows the water contents in the right ventricle, septum, and left ventricle of the control, continuous, and intermittent groups. The right ventricle and septum of each study group showed a slight, yet significant, increase in water content (right ventricle: continuous: +3.5% intermittent: +4.6%, p<0.002; septum: both groups: +3.5%, p<0.002) compared with that in the control group. No significant changes were observed in LV water content in the 3 groups.

**Table 2.** Myocardial water content.

| Cardiac Mass | Control | Continuous | Intermittent |
| --- | --- | --- | --- |
| Right ventricle | 78.4 ± 0.98 | 81.6 ± 0.37* | 82.5 ± 0.30* |
| Septum | 77.1 ± 0.85 | 79.8 ± 0.34* | 79.8 ± 0.22* |
| Left ventricle | 77.2 ± 0.77 | 79.1 ± 0.28 | 79.4 ± 0.26 |

n = 6 (control group); n = 7 (study groups); Values (%) = mean ± standard error;
\* p < 0.002 compared to respective control group values.

#### 3.3.3 VEGF EXPRESSION

Figure 4 shows the photomicrographs of the myocardium from animals of the three groups submitted to immunohistochemical staining for VEGF (Panels a, b and c). There are labeled cells (arrows) in the perivascular spaces, prominent and numerous in the intermittent group (Panel c). VEGF expression (% area fraction of immunolabeled histological sections) in the RV myocardium was significantly greater in the intermittent group (2.89% ± 0.41%) than in the continuous (1.80% ± 0.19%) and control (1.43% ± 0.18%) groups (p<0.023). VEGF expression in the myocardium of the right ventricle in the intermittent group was also greater than that in the left ventricle and septum within the same group (p<0.050; Figure 5). There was no significant difference in VEGF expression between the other cardiac sections or within the control and continuous groups.

**Figure 4.**
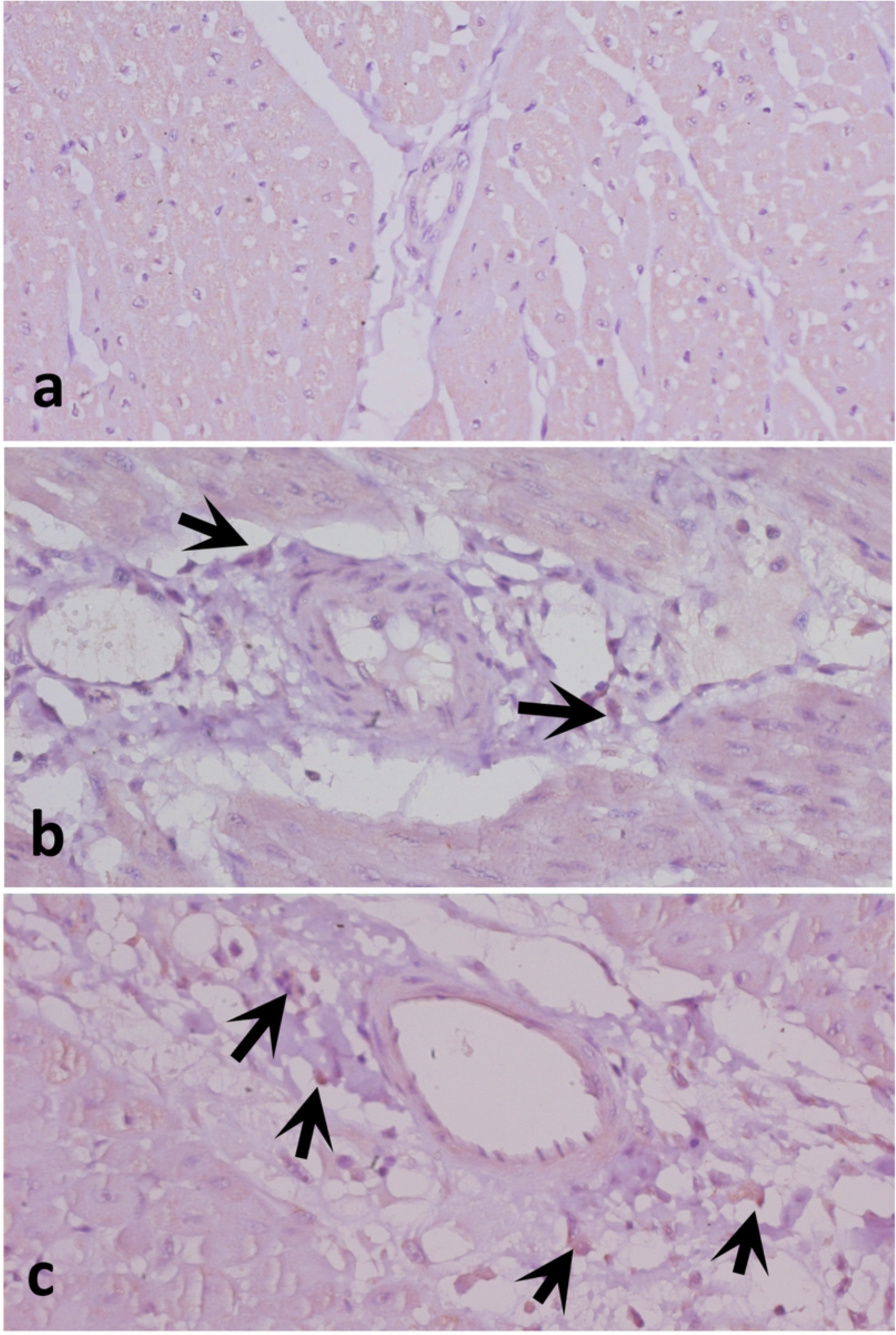
Photomicrographs of the myocardium from animals of the three groups (Panel a: Control; Panel b: Continuous; Panel c: Intermittent) submitted to immunohistochemical staining for vascular endothelial growth factor (VEGF). There are labeled cells (arrows) in the perivascular spaces, prominent and numerous in the intermittent group (Panel c).

**Figure 5.**
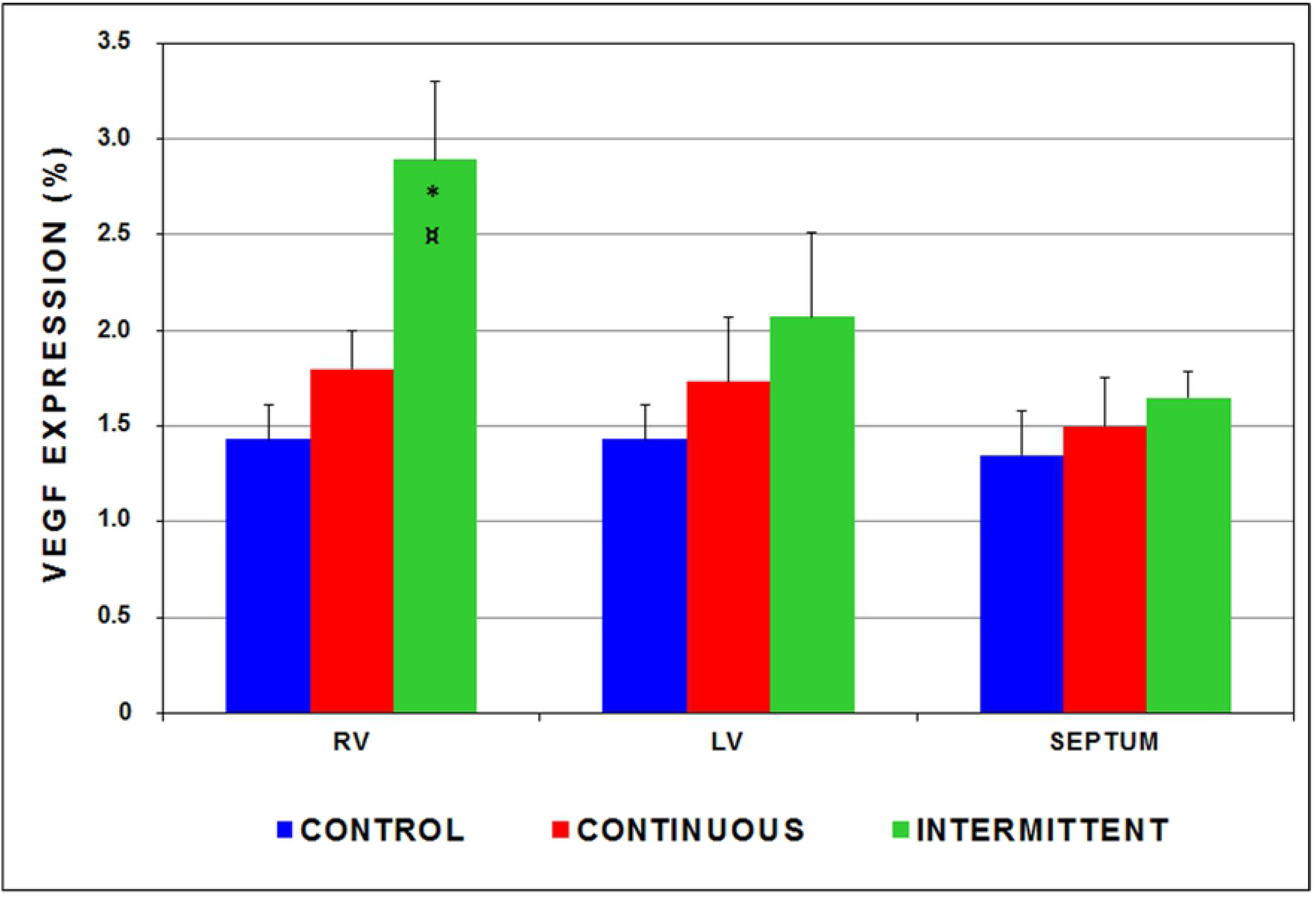
Vascular endothelial growth factor (VEGF) expression (% area fraction of immunolabeled histological sections) in the right ventricle (RV), left ventricle (LV), and septum in the 3 groups. * p<0.023 compared to RV of control and continuous groups; **¤** p<0.050 compared to LV myocardium and septum within the same group. n= 7 in each group.

## 4 DISCUSSION

This 96-hour protocol investigated the effects of subpulmonary ventricular retraining in young goats submitted to 2 programs of systolic overload on myocardial angiogenesis signaling via VEGF tissue expression. It has been clearly demonstrated that despite achieving a similar magnitude of RV hypertrophy in the 2 study groups, proportionally less exposure to systolic overload in the intermittent group promoted greater evidence of angiogenesis signaling than in the continuous group. Our research line has focused on the physiological hypertrophy associated with subpulmonary ventricular retraining, and we have found that intermittent systolic overload promotes myocardial hypertrophy without its undesirable effects. Therefore, morphologic assessment of adaptive mechanisms of the contractile and noncontractile myocardial elements submitted to the acute hypertrophy process can also significantly contribute to an understanding of the effects of ventricular retraining when the 2-stage Jatene operation is planned. The ultimate goal would be to minimize cell damage and maximize PAB efficiency, thereby, providing the best training program for the subpulmonary ventricle. The increased VEGF expression observed in animals from the intermittent group may represent one of the major physiological benefits to the myocardium. We believe that the RV myocardium, exposed to intermittent pressure overload and stretching of its myocardial fibers, has undergone adaptations at the cellular and molecular levels, with enhanced VEGF expression by endothelial and mesenchymal cells. Theoretically, such augmented expression should result in neovascularization in order to support the blood supply to the newly formed hypertrophic cardiomyocytes. The trigger for myocardial angiogenesis in the intermittent group might promote a healthier hypertrophy of the trained subpulmonary ventricle than traditional models of pressure-overload hypertrophy by increasing coronary vascular reserve and thus better energy supply. Previous studies have demonstrated a better myocardial performance index for the ventricle subjected to intermittent training. [9,17] Several mechanisms have been proposed to explain the better performance of the ventricle that is intermittently trained. VEGF is a well-documented, key regulator of vasculogenesis. [18] This key molecule participates in mechanisms that restore blood flow to tissues when the distribution of oxygen is impaired or the energy demand is increased. [19,20,21] According to cumulative knowledge gained in recent years, many transcription and additional growth factor molecules are involved in signaling pathways that regulate angiogenesis, such angiopioetin-1 and 2, fibroblast growth factor, transforming growth factor and platelet-derived growth factor. [22,23] VEGF and angiopoietins are, however, the prime regulators of myocardial angiogenesis. It is also recognized that, while during the adaptive phase of cardiac hypertrophy increased production of VEGF-A and angiopioietin maintain capillary density, inhibition of VEGF signaling result in capillary rarefaction and transition to heart failure. [24]

During ventricular retraining, the subendocardial pressure is expected to be increased by systolic overload imposed by PAB. Consequently, an imbalance with perfusion pressure and oxygen needs may promote subendocardial hypoxia. Hypoxia, in turn, is a powerful stimulus for vasculogenesis. Hypoxia leads to enhanced production of hypoxia-inducible factor-1 and the subsequent induction of VEGF and serum-derived factor-1. This signaling cascade stimulates endothelial cell progenitors during vasculogenesis. Endothelial cells differentiate from precursor cells, migrate, and form vascular channels. [25] This adaptive angiogenesis signaling is critical for perfusion in specific myocardial regions with limited coronary flow reserve, as in the subendocardium during subpulmonary ventricular retraining. On the other hand, it is possible that the complete relapse of the afterload during the rest condition may have facilitated tissue oxygenation and growth factor molecules synthesis when compared with the continuous group.

Also, recent experimental evidence links physiological cardiac hypertrophy secondary to exercise training with modulated expression of some microRNAs. [26] Expression of such microRNAs in paraffin-embedded tissue from our different study groups can give us a clue, in future analyses, of the quality of the adapted myocardium in our experimental model.

### 4.1 STUDY LIMITATIONS

The ventricular retraining protocol described here was carried out over a 96-hour period. Within this timeframe, significant changes in capillary density would not be expected to occur. Since VEGF signaling is one of the pathways for capillary proliferation, there should be a concordance of increased VEGF expression and a supposed increase in capillary density in the intermittent group. Nevertheless, an increase in VEGF signaling does not necessarily indicate capillary proliferation, since it is an oversimplification to assume and interpret the whole angiogenesis process based on the activity of a single molecule. As mentioned above, it is clear in the literature that other endothelial cell and tissue proliferation factors are related to myocardial angiogenesis. However, the limited commercial availability of primary antibodies against goat (the animal species of the present study) prevented us to broaden the search for other factors involved in vascular proliferation.

The expression of VEGF also has been studied in placental tissue, as this substance is related to the development of blood vessels responsible for fetal nutrition during pregnancy. [27] Even though placental tissue is highly vascularized, the relative expression of VEGF ranged only from 11% to 17% in uteroplacental tissue obtained from gilts at different gestational ages. This finding may help explain the low absolute levels of VEGF expression observed in our study, although altered degradation cannot be excluded. Although a relatively low percentage VEGF expression was observed in the goat myocardium, the finding of enhanced VEGF signaling in the intermittent group would be an early marker of sustained improved ventricular function during the 2-stage Jatene procedure. Nonetheless, it is important to emphasize that the changes have been measured over a 96-hour study period. One can argue whether the protocol had been extended, the changes in the 2 study groups would not have been similar.

### 4.2 CONCLUSIONS

This study demonstrates that intermittent versus continuous systolic overload promotes greater angiogenesis signaling. The association of RV hypertrophy with increased VEGF expression has important implications regarding the heart’s ability to adapt to pressure overload by promoting a compensatory growth of the coronary vascular reserve. Given that VEGF participates in mechanisms that restore blood flow to tissues when the energy demand is increased, we speculate that VEGF signaling can allow for a more efficient hypertrophy, and consequently, an optimized myocardial performance, a pivotal mechanism for the preservation of late ventricular function. Clinical trials of intermittent ventricular retraining for the 2-stage Jatene operation may translate these experimental findings into a long-term physiologic hypertrophy without late ventricular dysfunction.

## 6 ACKNOWLEDGEMENTS

Banding devices were provided by SILIMED Inc. (Rio de Janeiro, Brazil).

